# Chemical mapping exposes the importance of active site interactions in governing the temperature dependence of enzyme turnover

**DOI:** 10.1101/2021.06.25.449875

**Authors:** SD Winter, HBL Jones, DM Răsădean, RM Crean, MJ Danson, GD Pantoş, G Katona, E Prentice, VL Arcus, MW van der Kamp, CR. Pudney

## Abstract

Uncovering the role of global protein dynamics in enzyme turnover is needed to fully understand enzyme catalysis. Recently, we have demonstrated that the heat capacity of catalysis, Δ*C*_P_^‡^, can reveal links between the protein free energy landscape, global protein dynamics and enzyme turnover, suggesting that subtle changes molecular interactions at the active site can affect long range protein dynamics and link to enzyme temperature activity. Here we use a model promiscuous enzyme (Glucose dehydrogenase from *Sulfolobus Solfataricus*) to chemically map how individual substrate interactions affect the temperature dependence of enzyme activity and the network of motions throughout the protein. Utilizing a combination of kinetics, REES spectroscopy and computational simulation we explore the complex relationship between enzyme-substrate interactions and the global dynamics of the protein. We find that changes in Δ*C*_P_^‡^ and protein dynamics can be mapped to specific substrate-enzyme interactions. Our study reveals how subtle changes in substrate binding affect global changes in motion and flexibility extending throughout the protein.

---

A range of recent studies have begun to elucidate the relationship between global and local protein dynamics and enzyme turnover ^1–6^ There is now a range of computational and experimental evidence that variation in the normal distribution of vibrational modes, remote from the active site volume can affect the observed rate and temperature dependence of enzyme turnover. For example, evidence from simulation has demonstrated that differences in protein conformational sampling very distant from the active site volume can dramatically alter the temperature-dependence of catalysis.^7–8^ Similarly, isotope effect studies have pointed to significant changes in the thermodynamics of enzyme turnover, on small changes in vibrational frequency.^3, 9–12^ Indeed, there is some direct structural evidence for the proteins displaying long-range coherence of their vibrational modes, so called Fröhlich condensates.^13–14^ There is therefore building evidence for a complex relationship between local and long-range protein flexibility, catalysis and the temperature dependence of enzyme activity.

We have previously applied a model for understanding the temperature-dependence of enzyme catalysis (macromolecular rate theory, MMRT), which explicitly incorporates the difference in heat capacity between the ground state and transition state,

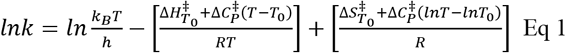

where T_0_ is an arbitrary reference temperature. 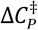 is the difference in heat capacity between the ground and transition state. 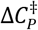 determines the change in Δ*H*^‡^ and Δ*S*^‡^ with temperature and thereby defines the non-linearity of the temperature dependence of the Gibbs free energy difference between the ground state and the transition state (ΔG^‡^).^15–17^ The magnitude of 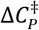 is a useful experimental window into the difference in rigidity between the ground state and the transitions state and more specifically the difference in the distribution of vibrational modes.

We have recently used a model hyperthermophilic enzyme system (a tetrameric glucose dehydrogenase from *Sulfolobus Solfataricus*; ssGDH) to study the contributions to the magnitude of 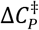.^1^ This experimental system has the advantage that it is extremely thermally stable and the chemical step (hydride transfer from a pyranose sugar to nicotinamide adenine dinucleotide phosphate [NAD(P)^+^]) is well defined and simple to monitor. ^18–19^ ssGDH is highly promiscuous and can turnover with a wide range of pyranose sugars as well as either NAD^+^ or NADP^+^, with only relatively small differences in *k*_cat_.^18–19^ In our recent study, we found an enormous kinetic isotope effect (KIE) on the magnitude of 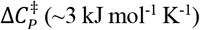, which is much larger than classical predictions would expect^1^. These data, combined with other recent efforts in the field, suggest a hypothesis where the vibrational modes of the protein are somehow coupled to the immediate active site volume, and influence the distribution of vibrational modes at either the ground or transition state.^20–21^

Herein we explore the hypothetical coupling of long-range protein vibrational modes to the active site volume, by removing individual hydrogen bonding interactions from the substrate sugar and monitoring the subsequent temperature dependence of enzyme turnover and changes in the network of enzyme dynamics through molecular dynamics simulation. Our approach therefore allows us to ‘chemically map’ the coupling of large-scale protein dynamics to the immediate active site volume and assess how this affects enzyme turnover. Moreover, using this approach, we illustrate the importance of the global protein dynamics in mediating the promiscuity of ssGDH by defining the distribution of active site conformational states. Our findings suggest that 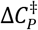 contributions can be dissected on the bond by bond level and not all the molecular interactions of the reactive complex geometry are necessarily pivotal for governing temperature dependence. Our data have implications for engineering enzyme substrate preference and the temperature dependence of enzyme activity.

## Results and Discussion

### Temperature dependence of enzyme kinetics with different substrates

We have previously monitored the temperature dependence of ssGDH turnover at relatively high temperatures in ssGDH (60-90 °C). We wished to explore whether this curvature was recapitulated across extremely long temperature ranges. To this end we have monitored the single-turnover kinetics of ssGDH with NADP^+^ and α-D-glucose across a 65 °C temperature range (20-85 °C) monitored as the change in absorption at 340 nm attributable to NADP^+^ reduction (Figure 1). Figure 1 shows the resulting temperature dependence of the observed rate, k_obs_. We found that at ‘low’ temperatures (20-50 °C) the data could be adequately fit to a sum of two exponential components (Figure 1, inset Eq), with the major phase at ~95% of the total amplitude (Figure 1A). At higher temperature (50-85 °C) we find the data are adequately fit to a single exponential function. Figure 1B, inset shows the resulting b factor from fitting to the exponential function (Figure 1A, inset Eq). This factor reflects the degree of ‘stretch’ of the exponential, with b = 1 reflecting an essentially ‘perfect’ exponential. Observing b ~ 1 suggests our data are not convolved of additional exponential phases other than those extract and deconvolved as above. Figure 1B shows the *k*_1_ values extracted from Eq 1 as a function of temperature. The curvature indicates a 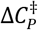 of −3.0 ± 0.6 for this expanded temperature range (20-85 °C), consistent (same within error) with our previously reported value for a smaller range^1^ (60-90 °C), −3.9 ± 1.1 kJ mol^−1^ K^−1^.

**Figure 1.**
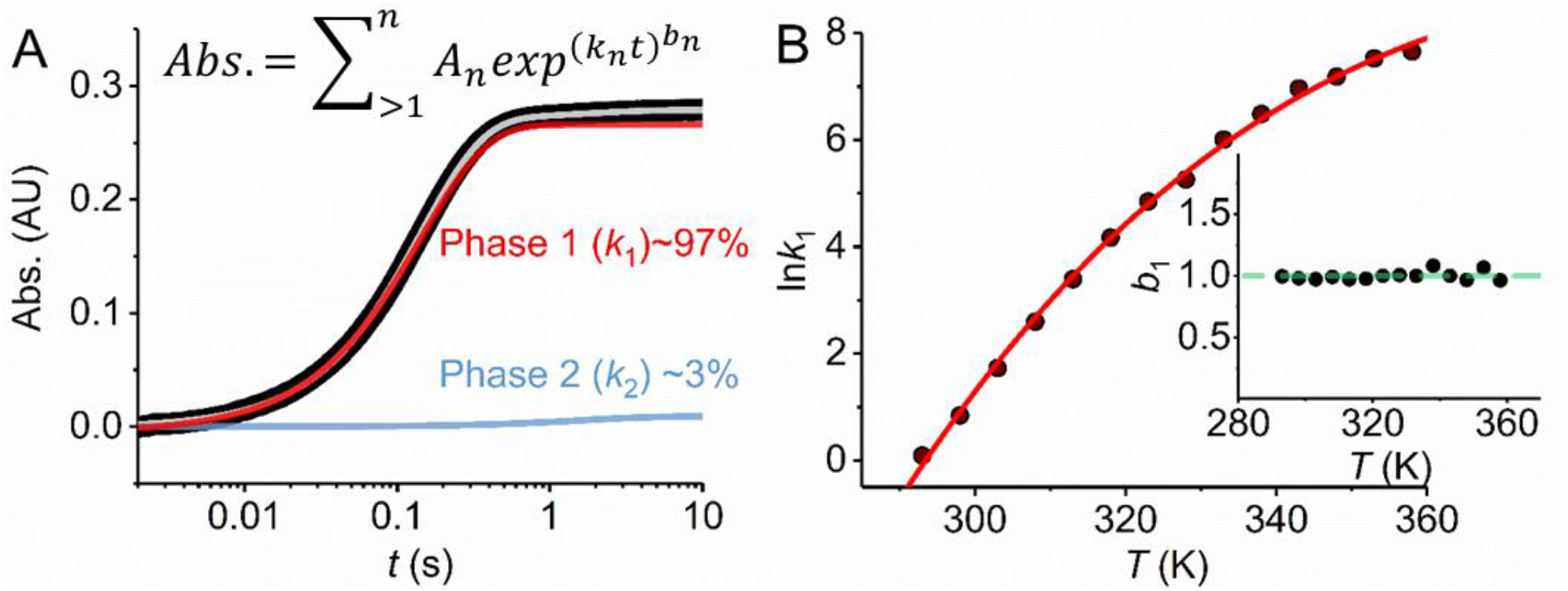
Long-range temperature dependence kinetics of GDH turnover monitored by stopped-flow. A, Example stopped-flow transient (black), fit using the inset equation (grey). B, Temperature dependence of the kinetic extracted from stopped-flow data. The solid red line is the fit to Eq 1. *Inset*, the temperature dependence of the magnitude of *b*, extracted from fitting to stopped-flow data as described in the main text. *Conditions:* 100 mM HEPES, pH 8.

We wish to explore how individual enzyme-substrate hydrogen bonding interactions affect the temperature dependence of ssGDH turnover. ssGDH is highly promiscuous and can accommodate a range of pyranose sugars. This combination of substrates therefore allows us to explore the sequential removal of hydrogen bonds arising from different hydroxyl groups on the sugar, either as a deoxy-monosaccharide or as the epimer as shown in Figure 2A and 2B. Using this approach, we are able to sensitively ‘chemically map’ the precise substrate bonding that governs the temperature dependence of enzyme turnover. Whilst single turnover measurements can be informative (Figure 1) we now turn to steady-state kinetics, which has the advantage of being more technically tractable and allowing us to explore the temperature dependence of *K*_M_ as well as to uncover more complex phenomena such as substrate inhibition if present. We show example Michaelis-Menten plots for each substrate used in Figure S1.

**Figure 2.**
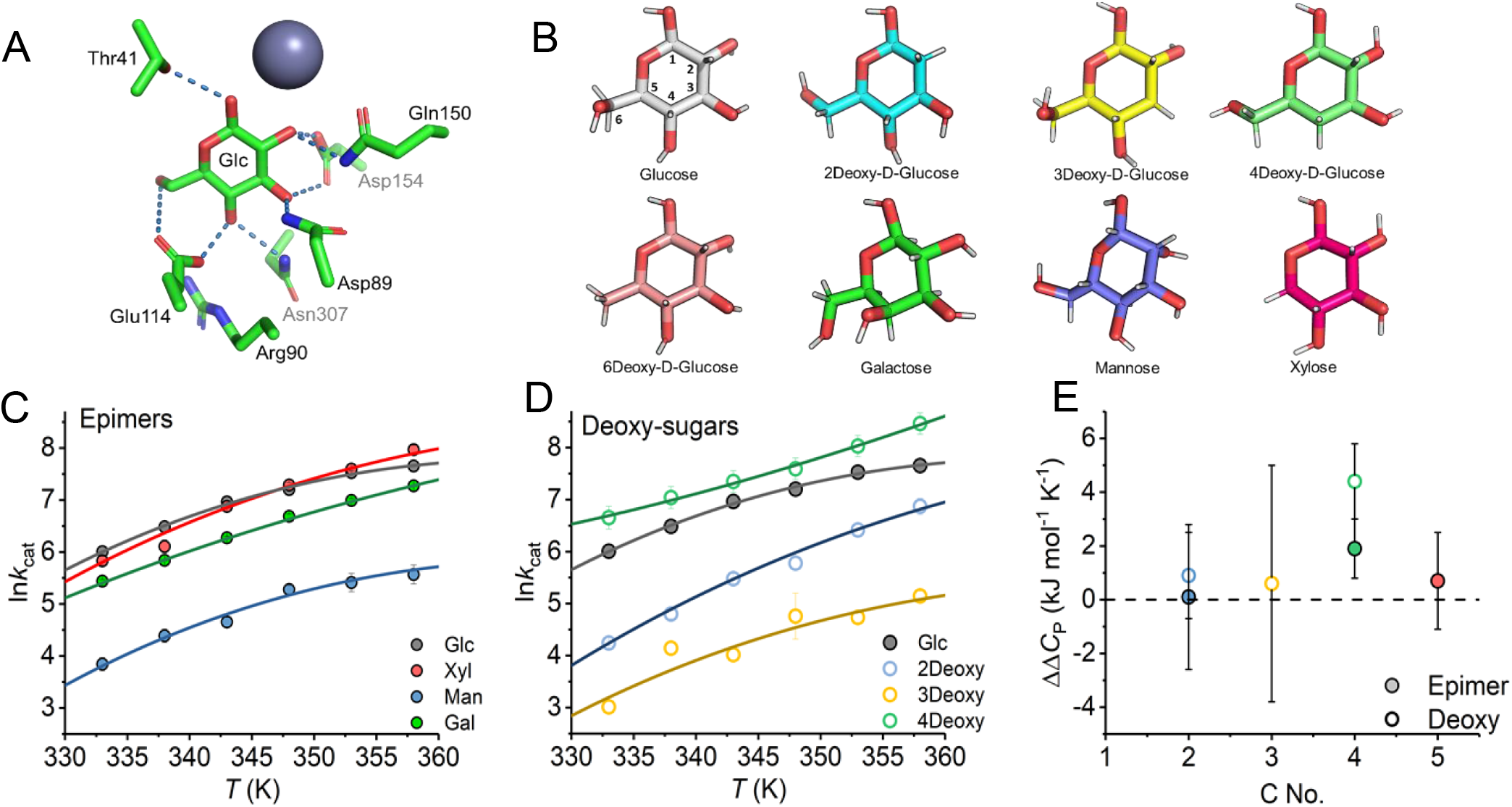
Chemically mapping the bonding contributions to the temperature dependence of ssGDH turnover. **A,** Key hydrogen bonding interactions between glucose and ssGDH. **B,** Glucose (with carbons numbered), its epimers and deoxy variants. **C, D,** Temperature dependence of *k*_cat_ for a range of substrates extracted from Michealis-Menten plots at each temperature (example plots shown in Figure S1). *Conditions,* 100 mM HEPES, pH 8. **E,** Change in magnitude of 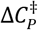 compared to D-glucose.

From the temperature dependence data (Figure 2C and 2D and Table 1) the magnitude of *k*_cat_ and *K*_M_ varies across ~ 2 orders of magnitude for different monosaccharides. In some cases, the *K*_M_ becomes rather large. Clearly one expects the binding component *K*_M_ to be affected on the loss of hydrogen bonds with each monosaccharide and so it is likely an increased dissociation constant accounts for a large part of the observed increase in *K*_M_. Notably, there is a trend for the variants at C2 (2-deoxy and D-mannose) to have very large *K*_M_ values whereas variants at C4 (4-deoxy-D-glucose and D-galactose) have *K*M values rather more similar to D-glucose. We note that we have not monitored the kinetics with D-allose because *ss*GDH essentially does not turnover with this monosaccharide, or at least is extremely slow relative to the other monosaccharides.^18^ We note that 4-deoxy-D-glucose shows apparent substrate inhibition (Figure S1) and this is not apparent for any of the other kinetic data. Despite the apparent substrate inhibition (*K*_i_ = 29.5 ± 8.7 mM), the extracted *k*_cat_ and *K*_M_ for 4-deoxy-D-glucose are similar to D-glucose with *k*_cat_/*K*_M_ being 1200 ± and 650 ± mM^−1^ min^−1^, respectively. Our molecular dynamics (MD) simulations below, provide a ready explanation for the apparent substrate inhibition, through the preference of the substrate for an inactive orientation and we discuss this in detail below.

**Table 1.** Steady-state and temperature dependence kinetic data for a range of substrate analogues.

|  | D-glucose<sup>b, c</sup> | 2-deoxy | 3-deoxy | 4-deoxy | D-xylose<sup>b</sup> | D-galactose | D-mannose |
| --- | --- | --- | --- | --- | --- | --- | --- |
| $k_{cat}$ (min<sup>-1</sup>)<sup>a</sup> | 651.9 ± 8.1 | 240.9 ± 8.0 | 60.9 ± 3.6 | 761.4 ± 12.8 | ~700 | 613.4 ± 21.2 | 141.1 ± 4.9 |
| $K_M$ (mM)<sup>a</sup> | 1.0 | 13.7 ± 3.7 | 56.6 ± 16.7 | 1.2 ± 0.1 | 0.5 ± 0.2 | 1.0 ± 0.1 | 75.5 ± 6.5 |
| $\Delta C_p^{\ddagger}$ (kJ mol<sup>-1</sup> K<sup>-1</sup>) | -3.0 ± 0.6 | -2.1 ± 1.0 | -2.4 ± 3.8 | 1.4 ± 0.8 | -2.3 ± 1.2 | -1.1 ± 0.4 | -2.9 ± 2.1 |
| $\Delta H^{\ddagger}$ (kJ mol<sup>-1</sup>)<sup>a</sup> | 63.6 ± 2.1 | 84.8 ± 4.2 | 73.0 ± 0.7 | 60.3 ± 10.0 | 80.4 ± 4.5 | 71.2 ± 1.4 | 56.3 ± 15.4 |

<sup>a</sup> data reported at 348 K. <sup>b</sup> values reported previously<sup>1</sup>. <sup>c</sup>Data from Figure 1.

We have previously interpreted changes in the magnitude of 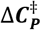 extracted from temperature-dependence plots to infer the presence of a difference in the distribution of vibrational modes between the ground and transition state^1, 22–24^. Figure 2E shows the change in the magnitude of 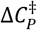 compared to D-glucose, 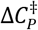, for each substrate. That is, 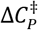 means no change in 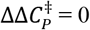 compared to D-glucose. From Figure 2C, 2D and Table 1, the absolute magnitude of 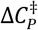 only varies (outside of experimental error) for both the C4-epimer (D-galactose) and the C4–deoxy substrate (4-deoxy-D-glucose). We note that the C4 position is distal from the immediate reaction centre (Figure 2A). Our data therefore suggest that the magnitude of 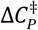 is highly sensitive to variation of the substrate at the C4 position, but no other individual position, and that this extends to both removal or inversion of the hydroxyl group.

We have previously solved the X-ray crystal structure of an inactivated variant of ssGDH with NADP and glucose/xylose bound.^19^ Here, we wished to explore the functionally active protein and so we have turned to molecular dynamics (MD) simulations to gain a structural understanding of the different potential bonding interactions of the reactive complex. Figure S2 shows the resultant conformations for a range of substrates by clustering on the root mean squared deviation (RMSD) values extracted from our MD simulations. The detailed results of the cluster analysis are shown in Figure S2. From the cluster analysis we find that the major enzyme-substrate conformations vary significantly between different substrates. Notably the 4-deoxy-D-glucose bound structure shows a significant rotation away from the catalytic zinc whereas the second highest orientation matches that of the rest of the sugars. Simulations with glucose and 6-deoxy-D-glucose both have one main conformation that makes up >80% of the population (Figure S2). These conformations occupy a similar orientation to one another. 2- and 3-deoxy glucose have a wide range of diverse conformations owing to the fact that these substrates have poor interactions with the protein and fall out of the active site in separate simulations after 60 to 80ns. This finding occurs in 4 of 20 cases (4 active sites from 5 independent simulations) for 3-Deoxy and 2 of 20 cases for 2-Deoxy.

The cluster analysis data are echoed in the calculated hydride donor-acceptor (D-A) distances (Figure S3) showing that the loss of specific H-bonding interactions give rise to a significant difference in the distribution of D-A distances with changes in both the range and magnitude of D-A distances. Indeed, 4-deoxy-D-glucose (Figure S3) populates a range of D-A distances, which are different from the other substrates, correlating with the clustering (Figure S2). Notably, unreactive distances are most frequently sampled. 4-deoxy-D-glucose also shows a very different distribution of hydrogen bond interactions (Table S3): Glu114 (instead of Thr41) now predominantly forms hydrogen bonds with the catalytically relevant O1 position. This change in substrate orientation, with a preference for large D-A distance, non-reactive geometries, is consistent with our observation of substrate inhibition with 4-deoxy-D-glucose. (We note that our previously reported D-A distances for glucose and xylose were based on less extensive conformational sampling^1^; our current results reveal a more complex D-A distance distribution). Our simulation data suggest that some of the observed differences in the *k*_cat_ might have an origin in altered D-A distances arising from changes in reactive complex geometry.

### Changes in protein dynamics with different substrates

We have previously found that the magnitude of 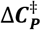 indicates the scale of the change in protein dynamics / distribution of vibrational modes between ground and transition state. The dynamical differences that give rise to altered 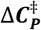 values can be as small as an isotopic substitution of the substrate^1^. Tracking relatively small changes in protein dynamics can be experientially challenging. We have recently developed the understanding of the protein red edge excitation shift (REES) phenomenon, specifically a quantification of the effect which allows subtle changes in protein dynamics to be tracked. ^25–28^ We term this quantification quantitative understanding of bimolecular edge shift, QUBES.^29^ We have demonstrated this quantification of the REES effect tracks subtle changes in protein dynamics, even where the crystal structures are identical and for a broad range of mutli-Trp proteins.^23, 29–30^ Briefly, REES is a fluorescence phenomenon where decreasing excitation energies are used to photo select for discrete emissive states, which manifests as inhomogeneous spectral broadening of the resultant emission spectra.^26, 31–32^

Using our QUBES approach we are able to interpret the REES effect to show relative changes in the equilibrium of discrete conformational sub-states. We wish to monitor differences in the flexibility of the enzyme ternary complex and so we have used a non-reactive mimic of NADPH, 1,4,5,6-tetrahydro NADP (NADPH_4_). We have elected to monitor the epimers using this approach because of their ready availability compared to the deoxy sugars. Figure 3A shows an example data set from which we track changes in the broadening of Trp emission spectra as the change in the center of spectral mass (CSM; See methods) with respect to excitation wavelength as in Figure 3B,

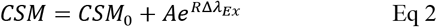

where the amplitude relative to *CSM*_0_ and curvature of the exponential is described by *A* and *R* values, respectively, and *CSM*_0_ is the CSM value independent of *λ_Ex_*. The *CSM*_0_ value reflects the solvent exposure of protein Trp residues manifest as a red shift in Trp emission on increasing solvent exposure.^33–35^ An invariant CSM_0_ value then suggests no significant structural change and this is the case for the present data set. We have previously found that, for an invariant CSM0 value, an increased A/*R* value is indicative of broader population of conformational states.^29^

**Figure 3.**
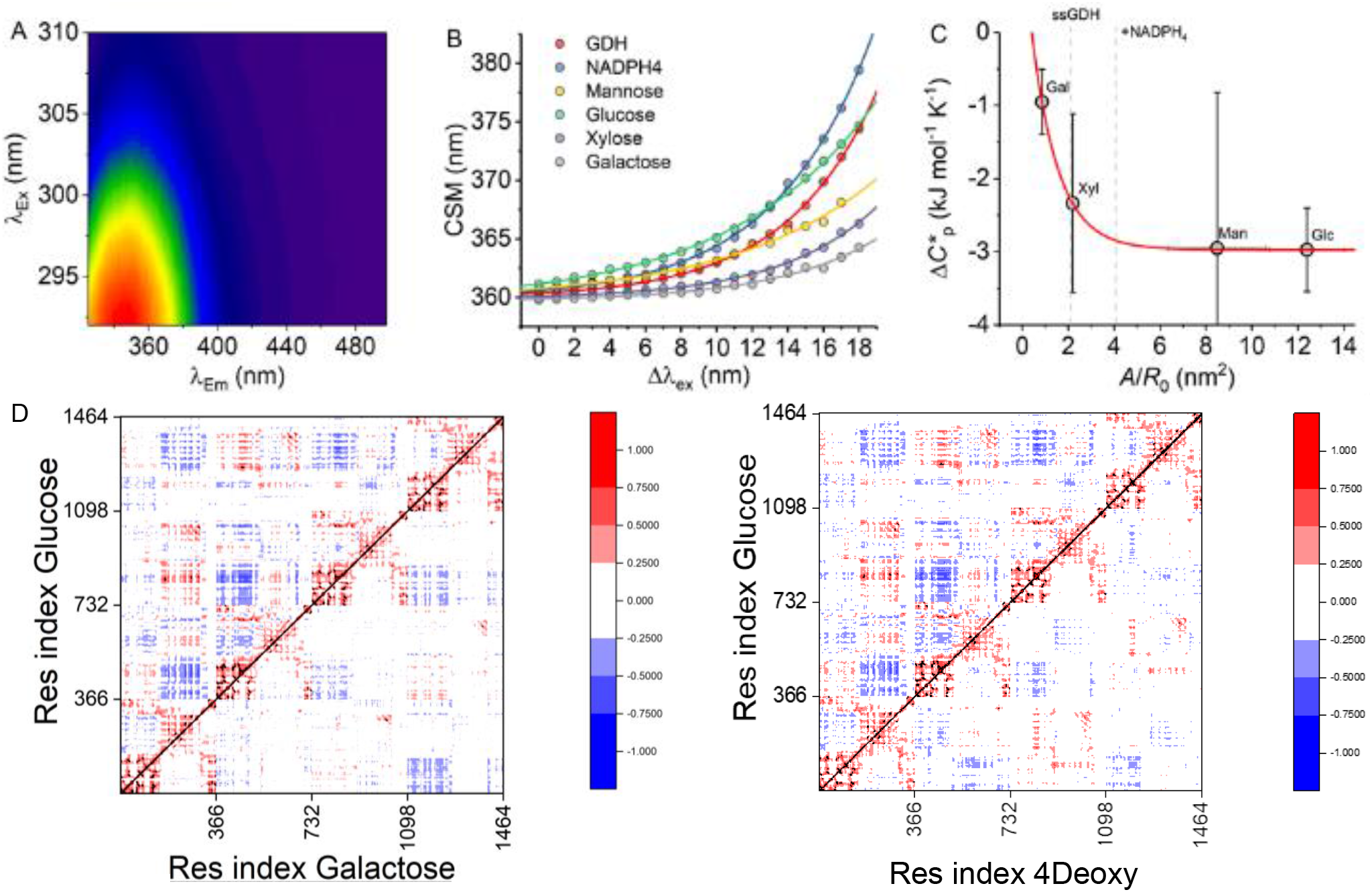
Changes in *ss*GDH flexibility and dynamics on substrate binding. **A,** Example raw REES data for *ss*GDH. **B,** Resulting CSM versus change in excitation wavelength extracted from panel A for a range of substrates + NADPH_4_. The solid line shows the fit to Eq 2, from which the QUBES data are extracted as the main text. **C,** Correlation between QUBES data and the magnitude of 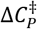 for different substrates. The solid red line is as simple exponential function fit to the data and is to aid the eye only. Grey dashed lines show the REES data for *ss*GDH and in the presence of NADPH_4_. Conditions: 100 mM HEPES, pH 8, 15uM GDH, 60 C°. **D,** Dynamic cross correlation for comparing glucose to Galactose and 4DGluc with a black diagonal line separating each system. Each new tick represents a new monomer. DCCM values are scaled between +1 (red, positively correlated motions between residues), 0 (white, no correlation) and −1 (blue, anti-correlated).

Figure 3C shows the resulting QUBES data for the ternary complex of GDH with a range of substrates *versus* the extracted magnitude of 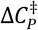 (Figure 2 and Table 1). Essentially, as the magnitude of *A*/*R* becomes larger the protein becomes more flexible; a broader range of accessible conformational substates as we have previously described.^23, 29–30^ From Figure 3C, we find that the REES data suggest that, despite the small number of H-bonding changes in the active site for each substrate, there is a significant difference in the equilibrium of conformational states with each of the epimers. We find that the ternary complex with D-galactose, outside of experimental error, explores a more restricted range of conformational states, i.e. is more ‘rigid’ compared to D-glucose. The data appear to show a trend of a decrease magnitude (increasingly negative values) of 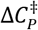 with a decreasing rigidity of the ssGDH ternary complex. We note that given the relatively large error associated with both the QUBES data and also the 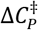, we can only confidently assert the difference between D-galactose and D-glucose. That is, at least for ssGDH a decrease in the flexibility of the enzyme ternary complex is associated with a decrease in the magnitude of 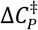.

The QUBES data provide a global metric of protein flexibility. To explore more localized contributions and to see if these are different/similar for the monosaccharides, we measure dynamical cross correlations and root mean square fluctuations (RMSF) from our MD simulations. Figure S4 shows per residue RMSF calculations for each of the substrates studied by MD. These data show a complex range of substrate specific changes in individual amino acids. There is not an obvious trend in the data that matches with the kinetic measurements. It is well established that in proteins, the motions of adjacent residues are often highly correlated with one another. This can produce a “domino effect” wherein perturbations to one residue create long-range interactions by propagating through networks of highly correlated neighbors. Dynamical cross-correlation matrices (DCCMs) can provide an image of how the protein interacts, not just within a monomer, but across the *ss*GDH tetramer. Specifically, DCCMs quantify the correlation coefficients of motions between atoms from which communication between distal residues can be inferred.^36^ We find a global loss in correlated and anticorrelated motions for all substrates in comparison with glucose (Figure 3D, Figure S5). Some of these differences are subtle, but the key finding is that the observed local changes in hydrogen bonding in the active site affect the protein dynamics across the tetrameric protein at large. D-galactose and 4-deoxy-D-glucose bound versions of GDH show a greater loss of correlated motion throughout the protein (in particular between the different monomers) compared to when glucose or other substrates are bound. This finding is intriguing, given our finding above that 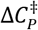 was markedly different for these substrates with modifications at the C4 position compared to other positions on the pyranose ring. Furthermore, our REES data indicate a that significant reduction in global protein flexibility when complexed with D-galactose.

## Conclusions

We have chemically mapped the specific bonding interactions that govern the temperature dependence of ssGDH turnover. We find a single set of bonding interactions is sufficient to drive a negative 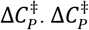 conceptually captures the difference in the distribution and frequency of vibrational modes at the ground and transitions state. Our REES data suggest the ground state ternary complex of ssGDH has a restricted range of conformational sub-states for D-galactose compared to D-glucose. Moreover, our molecular dynamics data imply that local changes in the hydrogen bonding network are sufficient to drive large scale changes in the correlation of protein motions throughout the tetramer. Taken together these data suggest that subtle changes in H-bonding pattern at the immediate active site volume not only translate to altered global protein dynamics, but also are sufficient to affect the distribution of vibrational modes at the ground and transition state. The extreme sensitivity of the functionally relevant protein dynamics mirror our previous findings showing the sensitivity to isotopic substitution and imply a fine balance between enzyme activity, temperature sensitivity and protein conformational dynamics.

## Methods

### ssGDH Expression and Purification

ssGDH was expressed with AmpR in a pET3a plasmid and grown on LB agar with ampicillin (100 μg/mL) at 37 °C. A 50 mL LB starter culture was used to inoculate 5 × 1 L LB until an OD_600_ of 0.5−0.6 was reached. Cells were harvested by centrifugation (4 °C, 14800 × g, 10 min) before being lysed by sonication using a lysis buffer (100 mM HEPES, pH 7.5), lysozyme, DNAase, and a protease inhibitor cocktail tablet. Soluble and insoluble fractions were separated by centrifugation at 4 °C (70000 × g, 10 min). Due to the thermostability of *ss*GDH, the soluble fraction was purified by heating the sample to 70 °C for 50 min. To remove precipitated protein, samples were centrifuged (4 °C 70000 × g, 10 min). The protein was further purified by passing the protein solution through a HiTrap Q HP anion exchange column. The protein was eluted over a gradient from 0 to 800 mM NaCl with 10 0mM HEPES (pH 7.5). The purification was completed using a HiLoad 16/600 Superdex column with an elution buffer 100 mM HEPES buffer (pH 8). Samples were dialyzed in 100 mM HEPES pH8 overnight. The ssGDH concentration was calculated spectroscopically using ε_260_ = 95 080 M^−1^ cm^−1^.

### Enzyme Assays

Stopped-flow measurements were conducted using a thermostated Hi-Tech Scientific stopped-flow apparatus (TgK Scientific, Bradford on Avon, UK). Typically 3 – 5 transients were recorded for each reported measurement. We used an excess of ssGDH in the presence of saturating concentrations of NADP^+^ and α-D-glucose as described in the main text, monitoring the change in absorption at 340 nm. Steady-state ssGDH kinetic measurements were carried out using a lidded 0.1 cm pathlength quartz cuvette and a UV−vis spectrophotometer (Agilent Cary 60 UV−vis spectrometer) in 100 mM HEPES (pH 8). Accurate concentrations of NADP^+^ were determined using the extinction coefficient of ε_260_ = 17 800 M^−1^ cm^−1^. Enzyme activity was measured for each condition at 85 °C by following the formation of NADPH at 340 nm using ε_340_ = 6220 M^−1^ as a direct measurement of ssGDH steady-state rates. Temperature dependent rates were calculated using concentration dependent data fitted to Michaelis Menten using *V*_max_. Kinetic data were gathered from 60 to 85 °C at 5 °C intervals. The data were fitted to eq 1 using OriginPro 2019b (MicroCal).

**Scheme 1.**
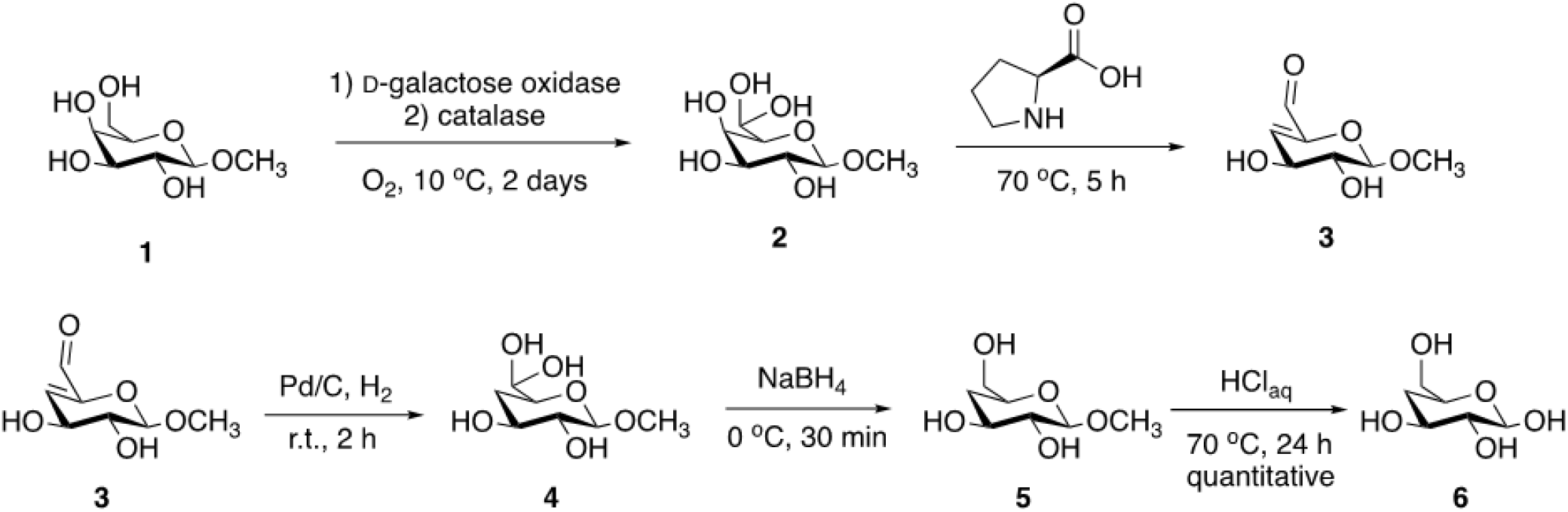
Synthetic route of 4-deoxy-D-glucose 6.

### Red Edge Excitation Shift Assays

for 5-10 minutes before measurements were performed at 60 °C. REES data was collected essentially as described previously as the matrix of excitation emission wavelengths.*^23, 29–30^* Fluorescence measurements were performed using a Perkin Elmer LS50B Luminescence Spectrometer (Perkin Elmer, Waltham, MA, USA) connected to a peltier heat pump (± 1 °C). 15 *μ*M ssGDH samples were used for analysis in 100 mM pH8 HEPES. NADPH_4_ was 100 μM where used. Samples were incubated Specifically, excitation was every nanometer between 292-310 nm, with emissions spectra collected between 325-500 nm. Excitation and emission slit widths were 4.5 nm in all cases. Temperature was regulated by a thermostated water bath at 60 °C. The CSM was extracted from the excitation emission matrix as,

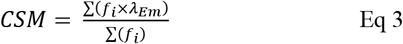

### 4-deoxy-D-glucose synthesis

13C NMR spectra were recorded on an Agilent Propulse instrument at 126 MHz frequency and 25 °C. Chemical shifts (δ) are reported in parts per million (ppm).

The synthetic route followed a modified version of a previously published chemoenzymatic procedure ^37^

#### Synthesis of 1 (as labelled in Scheme 1)

A 500 mL round-bottom flask was charged with methyl-β-D-galactopyranoside (500 mg, 2.58 mM, 1 equiv) dissolved in 130 mL 25 mM phosphate buffer (pH 7.3). The flask was sealed with a subaseal septum and cooled to 0 oC using an ice bath. Once cooled, oxygen was bubbled through the mixture for 5 min, followed by addition of galactose oxidase (2500 units) and catalase (20000 units). The flask was flushed with oxygen and the reaction mixture was stirred at 10 °C for 4 days. The product was used directly without isolation in the next steps.

#### One-pot cascade synthesis of 2 – 5

L-proline (30 mg, 0.26 mM, 0.1 equiv) was added to the solution of 1 and the reaction mixture was heated at 70 °C for 5 h to give 2. Once cooled to room temperature, a catalytic amount of palladium on activated carbon 10 % (10 mg, 0.09 mM, 0.03 equiv) was added and the mixture was stirred under hydrogen (atmospheric pressure) for 2 h. The catalyst was filtered off over Celite® and the filtrate was cooled to to 0 °C using an ice bath. To this, NaBH_4_ (49 mg, 1.29 mM, 0.5 equiv) was added in small portions over 30 minutes while keeping the solution cold. The unreacted NaBH4 was neutralised with acetone and the reaction mixture was passed through a glass column packed with mixed bed-resin TMD-8. To making sure all inorganic salts were removed, the filtrate was passed over TMD-8 several times until the resin changed colour. The solvent was removed under reduced pressure to yield product 5 as a white solid.

#### Synthesis of 6

Product 5 was resuspended in 2 M aqueous solution of HCl (pH 1.5) and the reaction mixture was heated at 70 °C for 24 h. The reaction progress was monitored by NMR and stopped when the peak corresponding to the methyl group disappeared. When conversion was complete, the reaction mixture was cooled to room temperature and the pH was adjusted to 7.3 with 0.1 M phosphate buffer. The solution was passed again through mixed bed-resin TMD-8 to remove the inorganic salts. The solvent was removed under pressure and the residue dried very thoroughly under high vacuum to yield 6 as a white sticky solid quantitatively. 13C NMR (126 MHz, D2O) δ: 103.8, 75.1, 72.7, 70.7, 68.6, 60.9, 57.1.

### Computational Methods

A previously prepared crystal structure 2CDB^1, 19^ was used as a starting model for each of the simulations. The Amber16 suite of programs was used for the periodic boundary simulation and analysis, with the ff14SB force-field for protein atoms, GLYCAM-06 for each of the sugars, TIP3P for water, parameters from Ryde and co-workers for NADP^+^ and ZAFF for the Zn^2+^ coordinated by Cys93, Cys96, Cys99 and Cys107. The catalytic Zn^2+^ was restrained to maintain the crystallographic coordination with Cys39 and His66. After brief minimization of the complex and added water, the system was heated to 300 K and subsequently equilibrated to 1 atm in the NPT ensemble (with positional restraints on Cα atoms). After gradual release of Cα positional restraints, 100 ns production simulations were performed at 300 K and 1 atm. Simulation analysis was performed using CPPTRAJ^38^. Hierarchical agglomerative clustering of substrate orientations was performed using the RMSD of non-hydrogen sugar atoms after alignment on the Ca atoms of active site residues (39, 41, 42, 66, 67, 89, 90, 105, 112, 114, 117, 150-151, 152-153, 192, 277, 279, 306-309, 313), with a minimum distance between clusters (epsilon) of 1.5. Root mean square fluctuation analysis were performed with CPPTRAJ using 10-100 ns of 5 independent simulations for each substrate. The data was averaged to give values for one monomer. The significant differences in RMSF values between no sugar bound and each of the other aforementioned sugar complexes were generated via a two-tailed T-test using R 3.4.3 software. Histograms of the D-A distances and determining of hydrogen bonding interactions between the sugars and protein and DCC analysis were calculated using CPPTRAJ as previously described for the RMSF data. DCC analysis was performed by generating an average RMSD of each of the α-carbons and measuring deviations from this model. Further details of model setup, restraints, and simulation procedures are included in the Supporting Information.

## ASSOCIATED CONTENT

### Supporting Information

Details of the experimental and computational approach, including simulation methodology RMSF and clustering analysis and associated supplementary figures and tables.

## AUTHOR INFORMATION

### Author Contributions

SDW, HBLJ and DR performed experimental work, all authors analyzed and interpreted data. All authors contributed to manuscript preparation. ǂThese authors contributed equally.

## ABBREVIATIONS

MMRT: Macromolecular Rate Theory
ssGDH: *Sulfolobus Solfataricus* glucose dehydrogenase
NADP: nicotinamide adenine dinucleotide phosphate
KIE: kinetic isotope effect
REES: red edge excitation shift
CSM: centre of spectral mass
DCCM: dynamic cross correlation mapping
RMSF: root mean square fluctuations.

---

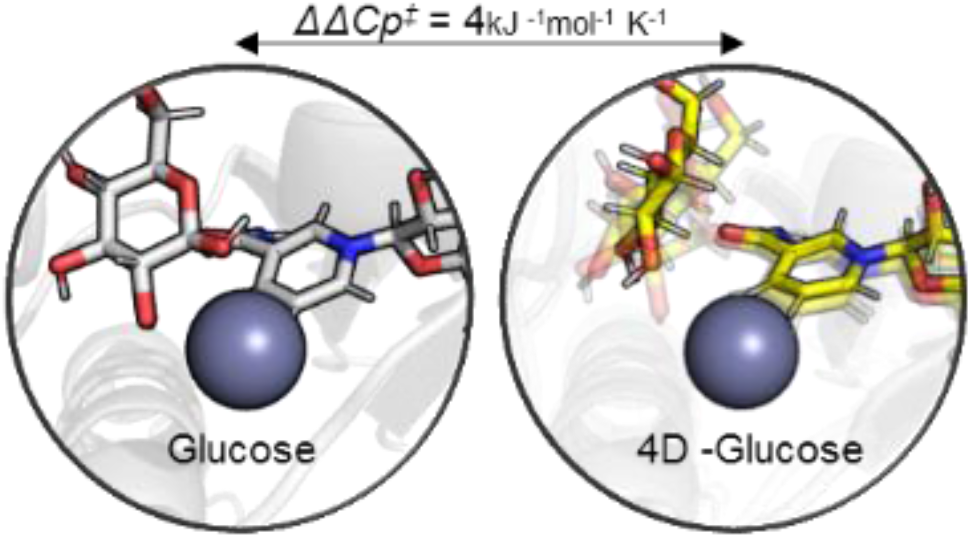
TOC

## Supplementary Computational Methods

### System Setup

As described in the main text, we previously performed MD simulations of glucose in complex with GDH,^1^ and we used this as the starting point for all simulations performed in this manuscript. This structure of glucose in complex with GDH was modified *in silico* as necessary to produce structures of GDH in complex with all of the sugars simulated herein. Protonation and histidine tautomerisation states, alongside any required Asn or Gln side chain “flips” were kept consistent for all systems and are the same as those used in our previous study.^1^ Histidine residues 66, 297 and 319 were singly protonated on their Nδ1 nitrogen, with all others singly protonated on their Nϵ2 nitrogen (as previously determined by REDUCE)^1^. All residues (bar the cysteine residues coordinating Zn^2+^) were simulated in their standard protonation states, consistent with our previously performed pKa calculations using PropKa 3.1.^39^ All structures were solvated in a rectangular water box (with all crystallographic water molecules retained) large enough such that no protein atom was within 10 Å of the box boundary. Na^+^ ions were added as required to make the total charge of the system neutral. The general equilibration procedure used for all MD simulations performed herein is described in the section titled “**Structure equilibration procedure**” below.

### Parameterisation of Modified Sugars

Complete GLYCAM-06j parameters for glucose, mannose and the 6-deoxy glucose variant are readily available from the original publication.^1^ For the 2,3 and 4-deoxy variants of glucose, all parameters bar the partial charges were available in GLYCAM-06j force field. Given the high similarity between glucose and each of its de-oxy variants, we made subtle chemically reasonable modifications to the partial charges of each sugar variant using the glucose partial charges as a template. (Only the carbon and its corresponding hydrogen’s charges at the position of the hydroxyl group removal were modified). These modified charges are provided in **Tables S1&S2**.

### Structure equilibration procedure

The following procedure was used to prepare all systems simulated for production MD simulations in the NPT ensemble at 300 K and 1 atm. All dynamics steps applied the SHAKE algorithm ^40^ to constrain all bonds containing hydrogen. Replicas simulations were initiated from the second heating step of the following protocol (with each replica therefore assigned different random velocity vectors at this stage). Simulations performed in the NVT ensemble used Langevin temperature control (with a collision frequency of 1 ps^−1^) and used a simulation timestep of 1 fs. Simulations performed in the NPT ensemble again used Langevin temperature control (collision frequency of 1 ps^−1^) and a Berendsen barostat (1 ps pressure relaxation time). The equilibration protocol is as follows: First, hydrogens atoms and solvent molecules were energy minimised (using 500 steps of steepest descent followed by 500 steps of conjugate gradient minimisation). To prevent the movement of non-hydrogen and non-solvent atoms during the minimisation, 10 kcal mol^−1^ Å^−1^ positional restraints were used to keep all heavy atoms fixed. Then the solvent was heated rapidly from 50 K to 300 K (NVT ensemble, 1 fs timestep) over the course of 200 ps, with the previously described restraints still maintained. The positional restraints were then replaced with 5 kcal mol^−1^ Å^−1^ positional restraints on only the Cα carbon atoms of each residue and subjected to another round of energy minimisation (500 steps of steepest descent followed by 500 steps of conjugate gradient). Retaining these positional restraints, the system was heated from 25 K to 300 K over the course of 50 ps (NVT ensemble, 1 fs time step). Simulations were then performed in the NPT ensemble (1 atm, 300 K, 2 fs time step) by first gradually reducing the 5 kcal mol^−1^ Å^−1^ Cα carbon restraints over the course of 50 ps. This was done by reducing the restraint weight by 1 kcal mol^−1^ Å^−1^ every 10 ps. A final 1 ns long MD simulation with no restraints on the Cα carbon atoms was then performed, with the final structure produced after this run, used as the starting point for production MD simulations. Please note that the restraints on the catalytic Zn^2+^ coordination sphere was used throughout the heating, equilibration and production simulations (and are described in the section titled: “**Restraints used During Production MD Simulations**”).

### Restraints used During Production MD Simulations

Consistent with our previously performed MD simulations,^1^ we applied three one sided harmonic restraints (commonly referred to as “wall potentials”) to the primary coordination sphere of the catalytic Zn^2+^ in order to maintain a catalytically competent position throughout the MD simulations. The three distance restraints were as follows: (1) His66 NE2−Zn^2+^ distances greater than 2 Å were restrained by a 70 kcal mol^−1^ Å^−2^ force constant. (2) Asp42 OD2−Cys39 N distances greater than 1.95 Å were restrained by a 70 kcal mol^−1^ Å^−2^ force constant. (3) Asp42 OD2−Zn^2+^ distances smaller than 4.2 Å were restrained by a 100 kcal mol^−1^ Å^−2^ force constant.

**Table S1.** Partial charges used to describe the 2-deoxy and 3-deoxy variants of glucose. Atom type labelling is consistent with standard GLYCAM-06j labelling.

| 2-deoxy Glucose (Residue Name: 2GB) |  |  | 3-deoxy Glucose (Residue Name: 3GB) |  |  |
| --- | --- | --- | --- | --- | --- |
| Atom Label | Atom Type | Partial Charge | Atom Label | Atom Type | Partial Charge |
| C1 | Cg | 0.384 | C1 | Cg | 0.384 |
| C2 | Cg | 0.029 | C2 | Cg | 0.31 |
| C3 | Cg | 0.284 | C3 | Cg | 0.007 |
| C4 | Cg | 0.276 | C4 | Cg | 0.276 |
| C5 | Cg | 0.225 | C5 | Cg | 0.225 |
| C6 | Cg | 0.282 | C6 | Cg | 0.282 |
| O1 | Oh | -0.639 | O1 | Oh | -0.639 |
| O3 | Oh | -0.709 | O2 | Oh | -0.718 |
| O4 | Oh | -0.714 | O4 | Oh | -0.714 |
| O5 | Os | -0.471 | O5 | Os | -0.471 |
| O6 | Oh | -0.688 | O6 | Oh | -0.688 |
| H1 | H2 | 0 | H1 | H2 | 0 |
| H1O | Ho | 0.445 | H1O | Ho | 0.445 |
| H21 | Hc | 0 | H2 | H1 | 0 |
| H22 | Hc | 0 | H2O | Ho | 0.437 |
| H3 | H1 | 0 | H31 | Hc | 0 |
| H3O | Ho | 0.432 | H32 | Hc | 0 |
| H4 | H1 | 0 | H4 | H1 | 0 |
| H4O | Ho | 0.44 | H4O | Ho | 0.44 |
| H5 | H1 | 0 | H5 | H1 | 0 |
| H61 | H1 | 0 | H61 | H1 | 0 |
| H62 | H1 | 0 | H62 | H1 | 0 |
| H6O | Ho | 0.424 | H6O | Ho | 0.424 |

**Table S2.** Partial charges used to describe the 4-deoxy variant of glucose. Atom type labelling is consistent with standard GLYCAM-06j labelling.

| 4-deoxy Glucose (Residue Name: 4GB) |  |  |
| --- | --- | --- |
| Atom Label | Atom Type | Partial Charge |
| C1 | Cg | 0.384 |
| C2 | Cg | 0.31 |
| C3 | Cg | 0.284 |
| C4 | Cg | 0.002 |
| C5 | Cg | 0.225 |
| C6 | Cg | 0.282 |
| O1 | Oh | -0.639 |
| O2 | Oh | -0.718 |
| O3 | Oh | -0.709 |
| O5 | Os | -0.471 |
| O6 | Oh | -0.688 |
| H1 | H2 | 0 |
| H2 | H1 | 0 |
| H3 | H1 | 0 |
| H41 | Hc | 0 |
| H42 | Hc | 0 |
| H5 | H1 | 0 |
| H61 | H1 | 0 |
| H62 | H1 | 0 |
| H1O | Ho | 0.445 |
| H2O | Ho | 0.437 |
| H3O | Ho | 0.432 |
| H6O | Ho | 0.424 |

**Table S3.**
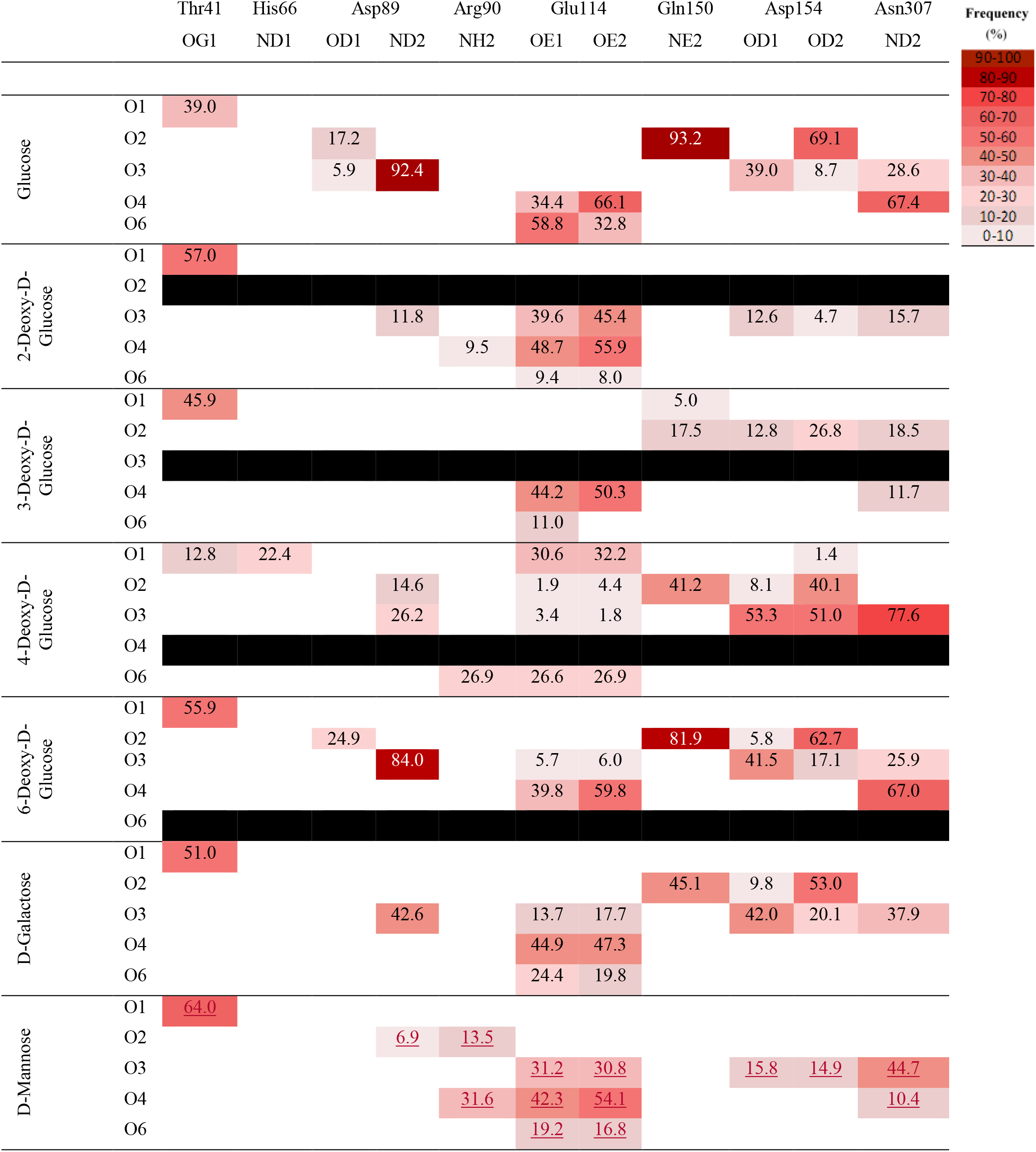
Hydrogen bonding network of ssGDH. The values represent the percentage of time over the simulation that there is a hydrogen bond from each hydroxyl of the sugar to each of the surrounding amino acids. Darker reds represent bonds which are present for longer periods of time.

**Figure S1.**
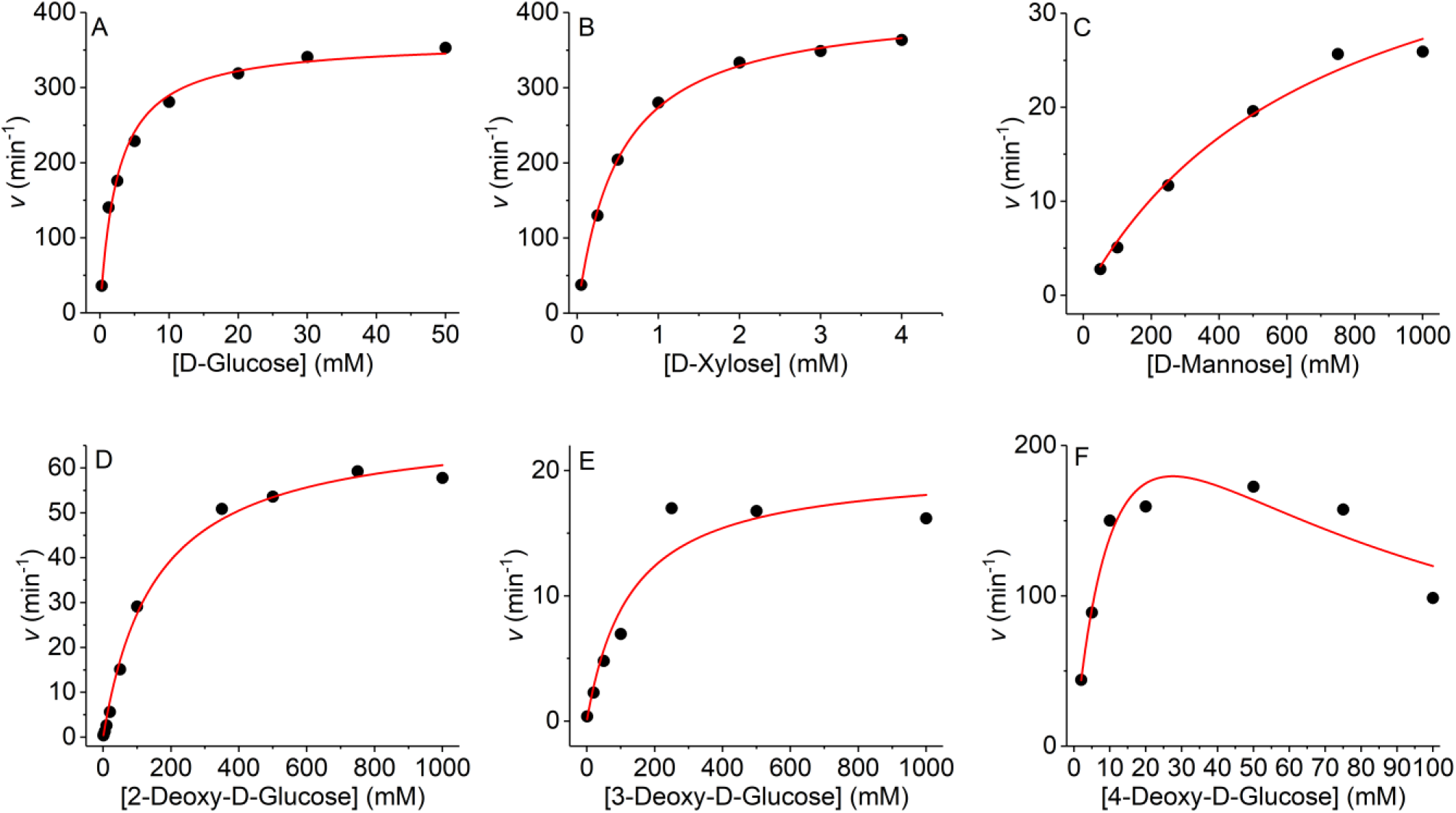
Example Michaelis-Menten plots for the substrates reported in the main manuscript with a saturating concentration of NADP (5mM) at 60 °C. The solid line is the fit to the Michaelis-Menten equation, 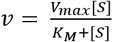 except in the case of 4-Deoxy-D-Glucose (panel F), which shows the fit to a function accounting for substrate inhibition, 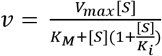.

**Figure S2.**
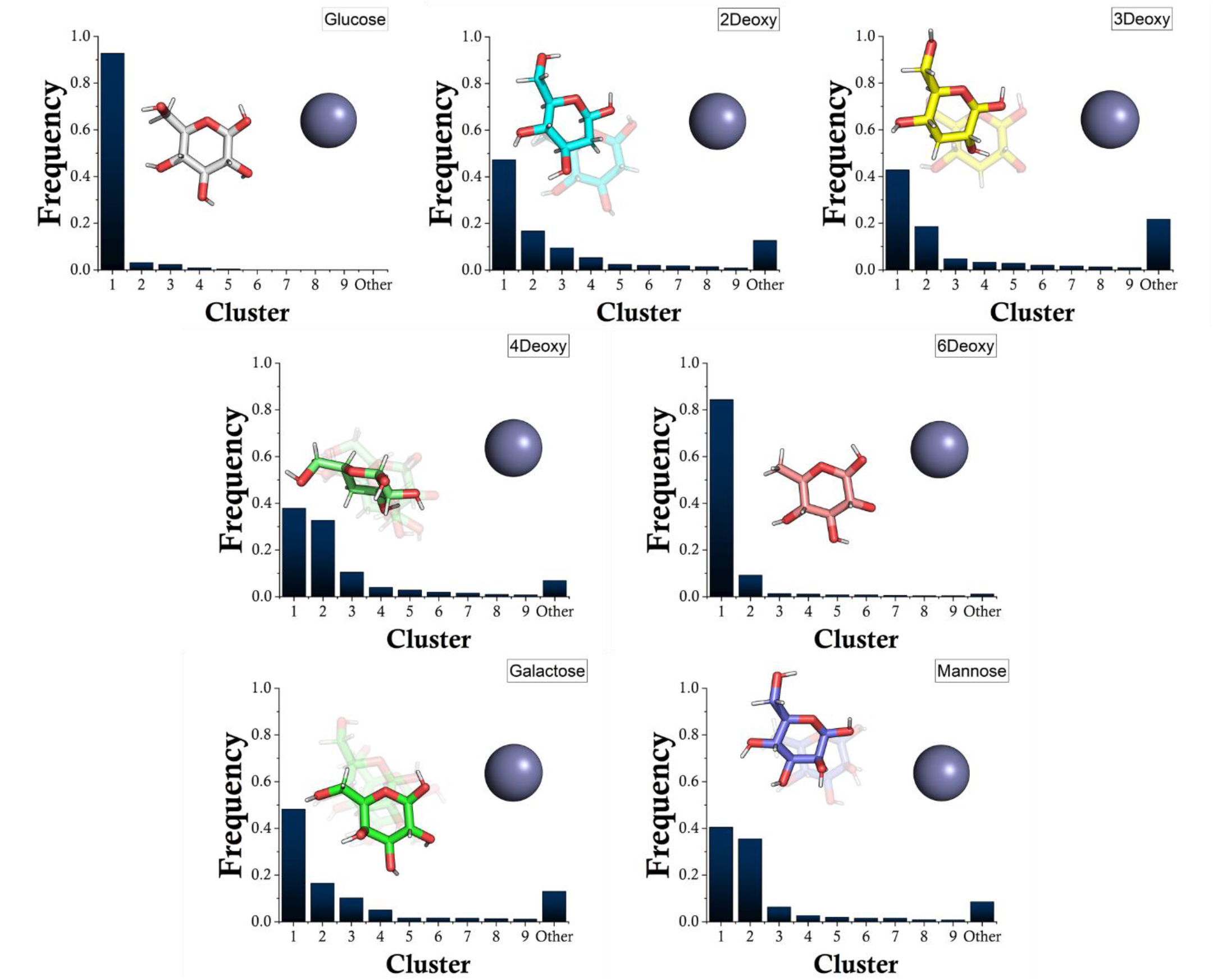
Different binding modes the contrasting sugars. The values represent the percentage of time over the simulation that the sugar exist within. Each of the images reflects the number 1 cluster i.e the highest occupied conformer and the transparent images represent lower frequency structures that exist with more than 0.1 frequency. Other refers to the percentage of omitted clusters form the simulation.

**Figure S3.**
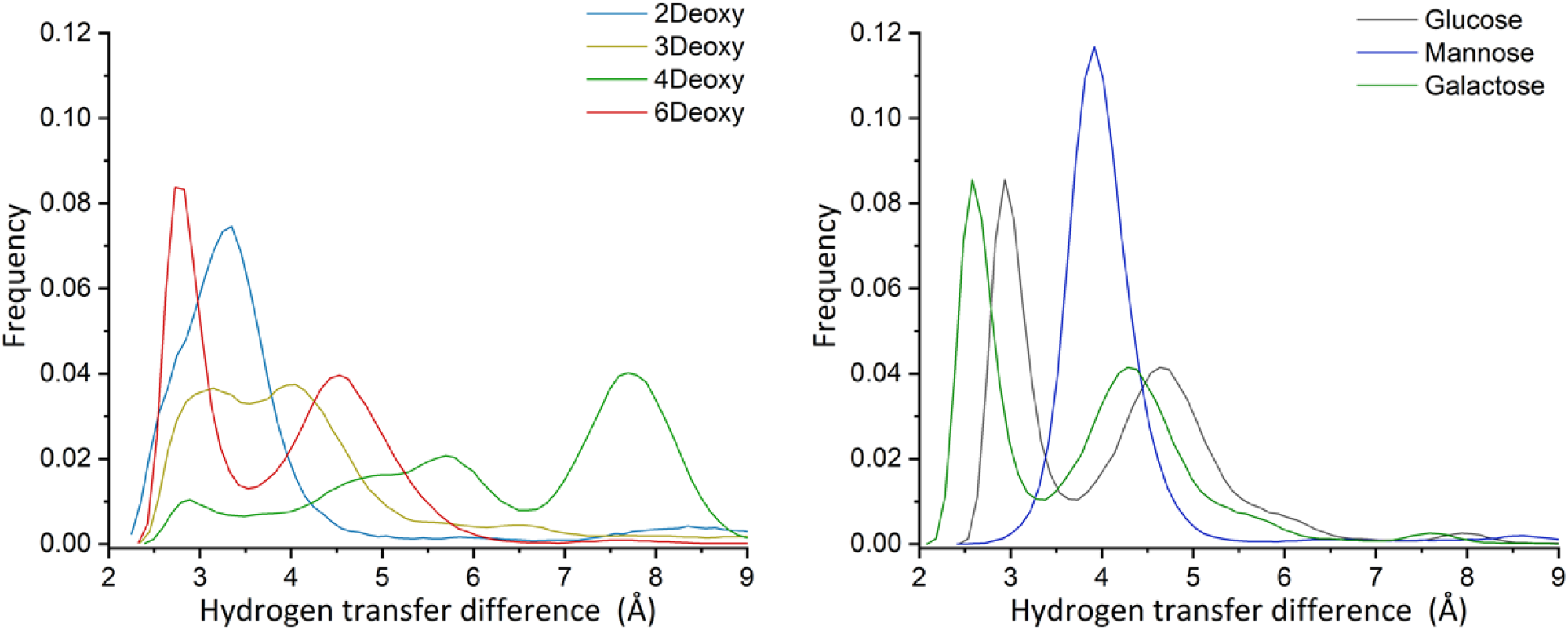
Histograms (bin width 0.1 Å) of the hydride donor-acceptor distances observed in MD simulations for each of the substrates (data from all 4 active sites in 5 independent simulations).

**Figure S4.**
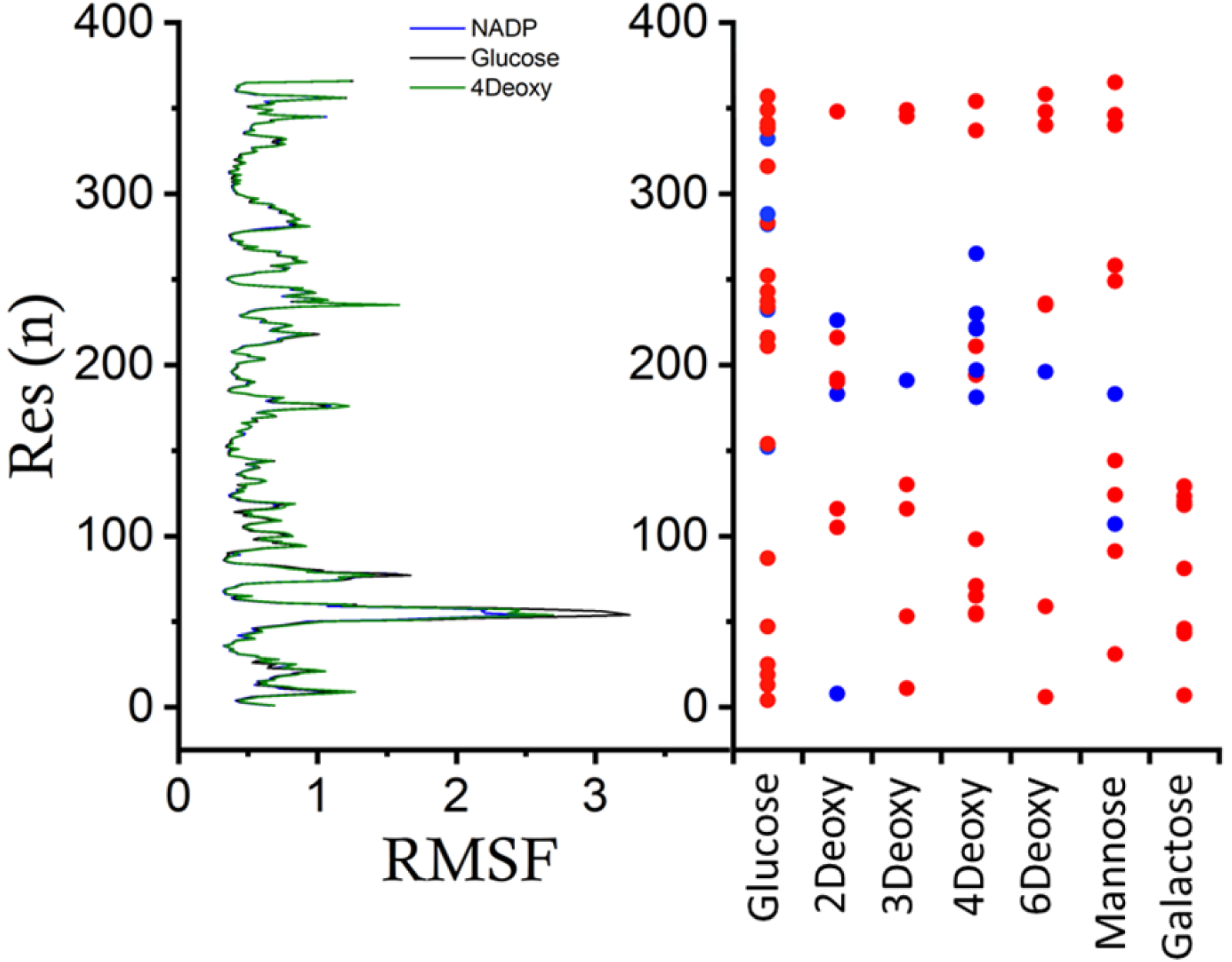
The Cα carbon root-mean square fluctuations per residue with no sugar bound together with residues that change significantly in fluctuation for the simulations with sugar bound. Significant differences in fluctuation vs. simulation without sugar are determined using a two-sided t-test based on the RMSF of the 20 protein chains for each complex (4 from each of the 5 independent simulations per complex). Red symbols signify residues that are significantly more flexible, blue symbols signify residues that are significantly less flexible.

**Figure S5.**
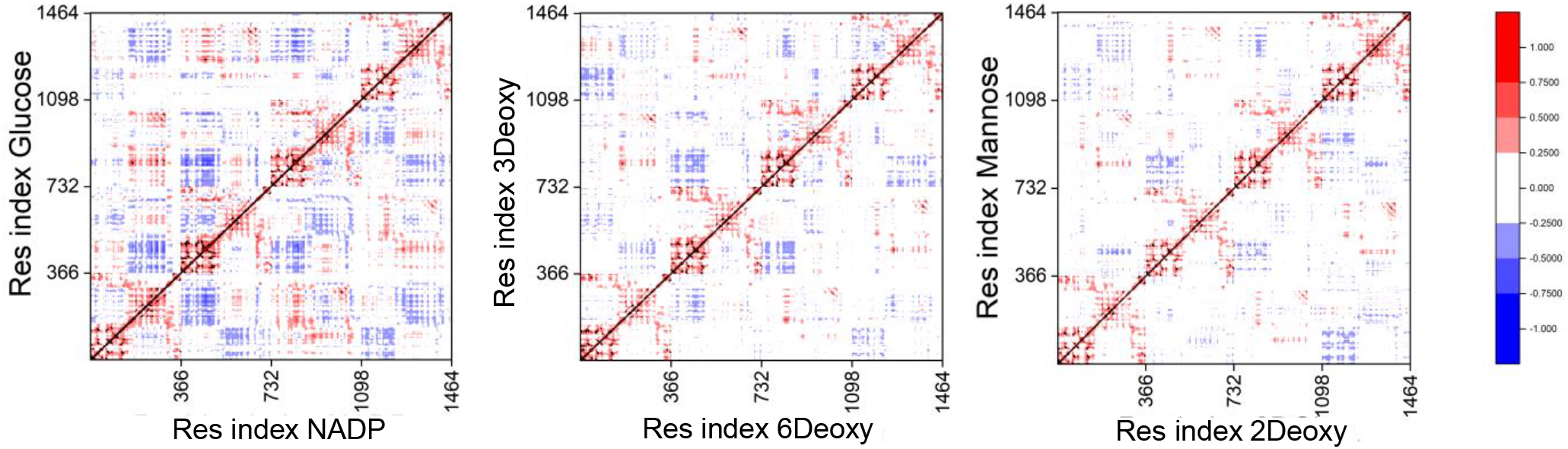
Dynamic Cross Correlation matrices. Black diagonal line separates each system. Each new tick represents a new monomer. DCCM values are scaled between +1 (red, positively correlated motions between residues), 0 (white, no correlation) and −1 (blue, anti-correlated).

